# Scavenger receptor class B type I is required for efficient glucose uptake and metabolic homeostasis in adipocytes

**DOI:** 10.1101/2023.08.21.554190

**Authors:** Darcy A. Knaack, Jackie Chang, Michael J. Thomas, Mary G. Sorci-Thomas, Yiliang Chen, Daisy Sahoo

## Abstract

Obesity is a worldwide epidemic and places individuals at a higher risk for developing comorbidities that include cardiovascular disease and type 2 diabetes. Adipose tissue contains adipocytes that are responsible for lipid metabolism and reducing misdirected lipid storage. Adipocytes facilitate this process through insulin-mediated uptake of glucose and its subsequent metabolism into triglycerides for storage. During obesity, adipocytes become insulin resistant and have a reduced ability to mediate glucose import, thus resulting in whole-body metabolic dysfunction. Scavenger receptor class B type I (SR-BI) has been implicated in glucose uptake in skeletal muscle and adipocytes via its native ligands, apolipoprotein A-1 and high-density lipoproteins. Further, SR-BI translocation to the cell surface in adipocytes is sensitive to insulin stimulation. Using adipocytes differentiated from ear mesenchymal stem cells isolated from wild-type and SR-BI knockout (SR-BI^-/-^) mice as our model system, we tested the hypothesis that SR-BI is required for insulin-mediated glucose uptake and regulation of energy balance in adipocytes. We demonstrated that loss of SR-BI in adipocytes resulted in inefficient glucose uptake regardless of cell surface expression levels of glucose transporter 4 compared to WT adipocytes. We also observed reduced glycolytic capacity, increased lipid biosynthesis, and dysregulated expression of lipid metabolism genes in SR-BI^-/-^ adipocytes compared to WT adipocytes. These results partially support our hypothesis and suggest a novel role for SR-BI in glucose uptake and metabolic homeostasis in adipocytes.

## Introduction

Over the last several decades, obesity has nearly tripled in adults^1,2^ and children^2,3^ and continues to be a worldwide epidemic. Obese individuals are at a higher risk than non-obese individuals for developing comorbidities, including cardiovascular disease, type 2 diabetes, some cancers, and premature death (reviewed in^4^). Within adipose tissue, adipocytes store dietary metabolites (i.e., free fatty acids, glucose) in the form of triglycerides to avoid misdirected lipid storage within the body. These triglycerides can then be catabolized to meet our body’s energetic demands. Although storing fat within adipocytes serves as a protective mechanism, the increase in Westernized diet consumption of processed fats and sugars has led to excessive fat accumulation and increased inflammation, thereby causing adipocytes to become dysfunctional and metabolically imbalanced^5^. An underappreciated role of adipose tissue is its ability to function as an endocrine organ that signals and communicates with other organs by secreting hormones and adipokines (i.e., adiponectin, leptin) to maintain whole-body metabolic homeostasis^6,7^. As such, the disruption of metabolic function in adipose tissue could lead to whole-body metabolic dysfunction, otherwise known as metabolic syndrome. Thus, elucidating the cellular mechanisms that may contribute to metabolic imbalance within adipose tissue has become an increasingly attractive area of research to better understand the forces that drive the development of obesity, metabolic syndrome, and other comorbidities.

Apart from their role in lipid storage, adipocytes are largely responsible for facilitating glucose metabolism. In the fed-state, the pancreas of a normal, healthy individual releases insulin into circulation. Once insulin binds to the adipocyte insulin receptor, downstream signaling through the PI3K/AKT pathway stimulates the translocation of insulin-sensitive glucose transporter 4 (GLUT4) from intracellular storage vesicles to the plasma membrane^8^. GLUT4 then facilitates the transport of glucose molecules into the adipocyte, which can be metabolized to synthesize triglycerides for storage or provision of cellular energy as adenosine triphosphate (ATP) through glycolysis or the electron transport chain (reviewed in^9^). Under obese conditions, adipocyte function is hindered due to the loss of insulin responsiveness (reviewed in^10^). These circumstances lead to insulin being unable to inhibit lipolysis, or the breakdown of triglycerides into fatty acids, thus leading to increased release and accumulation of fatty acids in circulation^11^. The resulting adipocyte insulin resistance and misdirected lipid storage within the body calls out the need to better understand the role of adipocyte proteins that enable the import and/or processing of metabolic intermediates that contribute to obesity and its comorbidities.

Scavenger receptor class B type I (SR-BI) is a class B transmembrane scavenger receptor that is primarily expressed in the liver, steroidogenic tissues, and adipose tissue^12–14^. Serving as the primary receptor for high-density lipoproteins (HDL)^15^, SR-BI’s role in cholesterol transport has been extensively studied in diverse tissue types (reviewed in^16^). As adipocytes have low *de novo* cholesterol biosynthesis^17^, they rely on lipoproteins for their cholesterol supply^18,19^. In adipocytes, SR-BI facilitates bidirectional cholesterol transport *in vivo* by the efflux of free cholesterol to HDL particles^20^, as well as by the selective uptake of cholesteryl esters from HDL^21^. In addition to facilitating cholesterol transport, SR-BI performs other functions within adipose tissue, including fatty acid transport^22^ and glucose uptake^23^. In the latter study, Zhang et al.^23^ demonstrated a role for HDL in stimulating glucose uptake into 3T3-L1 adipocytes by SR-BI^22^. Considering that SR-BI translocation to the cell surface in adipocytes is sensitive to insulin stimulation^24^, we propose that SR-BI might also be important for insulin-stimulated glucose uptake. In this study, we use adipocytes differentiated from ear mesenchymal stem cells (EMSCs) isolated from genetically modified mouse models to test the hypothesis that SR-BI is required for insulin-mediated glucose uptake and regulation of energy balance in adipocytes.

## Results

### Mesenchymal stem cells isolated from ears of WT and SR-BI^-/-^ mice differentiate into mature adipocyte-like cells

Hallmark characteristics of *in vitro* stem cells differentiated into adipocytes include increased neutral lipid accumulation within lipid droplets, induction and expression of adipogenesis genes and proteins, as well as secretion of adipokines (e.g., adiponectin). To verify adipogenesis, EMSCs isolated from wild-type (WT) and SR-BI knockout (SR-BI^-/-^) mice were cultured to passage 3 and differentiated over a 9-day period. Cells at days 0, 3, 5, 7, and 9 post-differentiation were stained with Oil Red O (ORO) to assess neutral lipid accumulation, imaged (**Figure 1A**), and quantified by isopropanol extraction (**Figure 1B**). Both genotypes displayed lipid droplet maturation and increased levels of neutral lipid accumulation over the post-differentiation period compared to undifferentiated cells (day 0), with adipocytes at days 7 and 9 post-differentiation showing the greatest amount of staining. As lipid accumulation seemed to plateau by day 9 post-differentiation, we considered adipocytes between days 7 and 9 post-differentiation as mature adipocytes. We further validated this model system through passage 5 by ORO imaging and quantification (**Supplemental Figure 1**). However, the capacity for these cells to differentiate into adipocytes appears to decrease with increasing passage numbers (**Figure 1** and **Supplemental Figure 1**) and may account for the variability observed within our study, thus leading us to present normalized data versus raw data. As such, all experiments in this study were performed in adipocytes between days 7 and 9 post-differentiation and between passages 3 and 5, unless otherwise stated.

**Figure 1.**
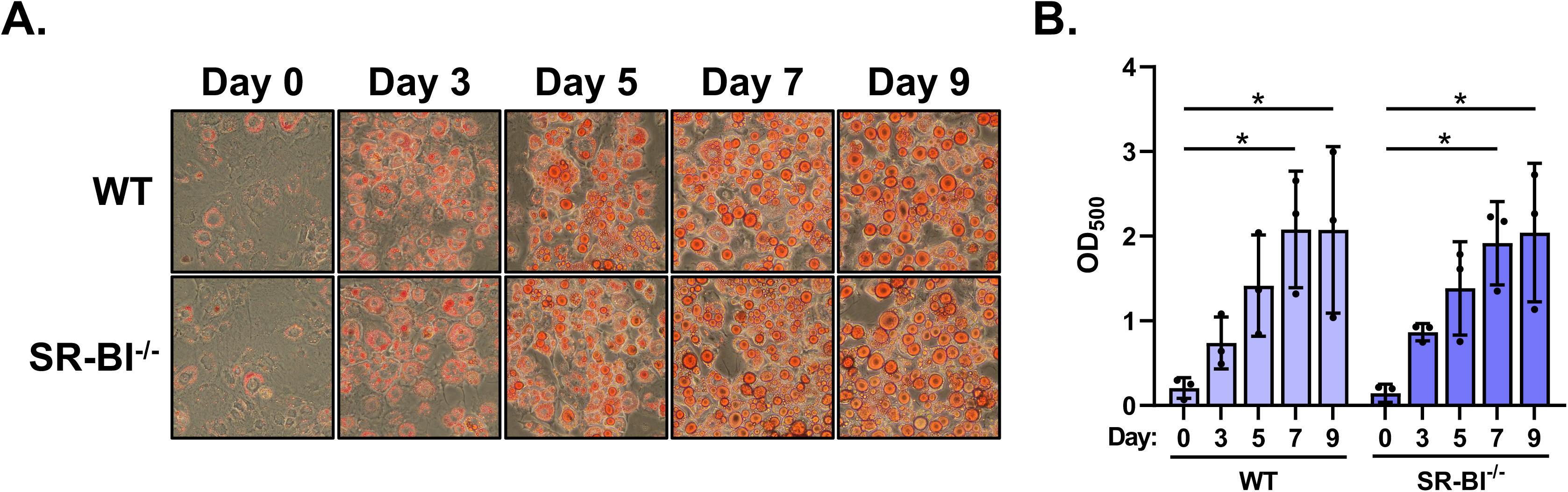
EMSCs isolated from WT and SR-BI^-/-^ mice differentiate into adipocyte-like cells. **(A)** EMSCs from WT and SR-BI^-/-^ mice at days 0, 3, 5, 7, and 9 post-differentiation were stained with ORO and imaged at 20X magnification. Representative images were selected (n=3 images/day/genotype) from n=3 independent EMSC isolations. **(B)** ORO staining was quantified by extracting the dye using isopropanol and absorbance at 500 nm was measured. Data are presented as mean ± SD (n=3, two-way ANOVA, Tukey’s post hoc [*p≤0.05]).

We next analyzed whether adipocytes differentiated from WT and SR-BI^-/-^ EMSCs displayed upregulation of adipose-specific genes over the post-differentiation period. Compared to undifferentiated EMSCs at day 0, WT and SR-BI^-/-^ adipocytes at days 7 and 9 post-differentiation showed the expected upregulation of *Cebpa* (CCAAT/enhancer-binding protein [C/EBPɑ]) and *Pparg2* (peroxisome proliferator-activator receptor γ [PPARγ] isoform 2 [PPARγ2]) (**Figures 2A** and **2B**). We also measured gene expression of *Pref1* (preadipocyte factor 1 [Pref-1]), which is mainly expressed in preadipocytes^25^. Compared to undifferentiated EMSCs at day 0, WT and SR-BI^-/-^ adipocytes at days 7 and 9 post-differentiation showed downregulation of *Pref1* (**Figures 2A** and **2B**). Protein expression of perilipin-1, a lipid droplet-associated protein, and PPARγ2 in whole-cell lysates was determined by immunoblot analysis (**Figure 2C**). We observed that WT and SR-BI^-/-^ adipocytes had induced expression of perilipin-1 and PPARγ2 at days 7 and 9 post-differentiation compared to undifferentiated cells at day 0. The antibody used to detect PPARγ recognized both of its isoforms, PPARγ1 and PPARγ2. The PPARγ2 isoform is induced during adipocyte differentiation and expression is specific to adipose tissue, whereas the PPARγ1 isoform is more ubiquitously expressed^26^. Interestingly, we observed that SR-BI^-/-^ EMSCs had decreased expression of *Pref1* and *Cebpa*, and increased expression of *Pparg2* in undifferentiated cells (day 0) compared to WT EMSCs (**Supplemental Figure 2**). These data suggest that SR-BI may be important for the adipocyte differentiation process and that transcriptional upregulation of *Pparg2* may be independent of *Cebpa* expression in adipocytes.

**Figure 2.**
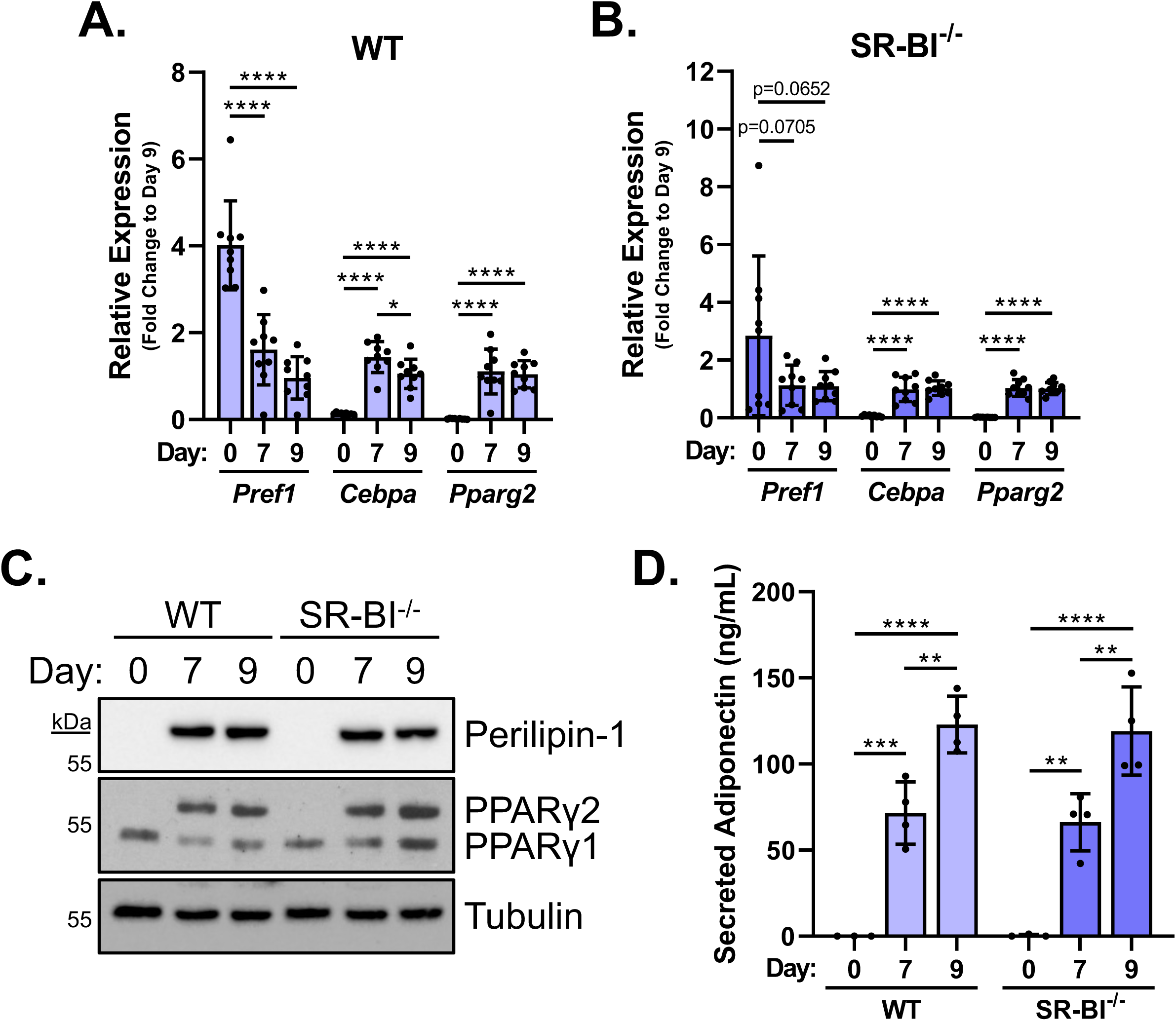
EMSCs differentiated into adipocytes from WT and SR-BI^-/-^ mice express markers of adipogenesis. Gene expression of preadipocyte marker *Pref1* (Pref-1) and adipocyte markers, *Cebpa* (C/EBPα) and *Pparg2* (PPARγ2) was assessed over the post-differentiation period in **(A)** WT and **(B)** SR-BI^-/-^ adipocytes by the 2^-ΔΔCt^ method using *Rplp0* (60S ribosomal subunit) as a housekeeping gene. Data were then subsequently normalized as a fold change to day 9 post-differentiation and presented as mean ± SD (n=3 independent experiments performed in duplicate or triplicate, one-way ANOVA, Tukey’s post hoc, [*p≤0.05, ****p≤0.0001]). **(C)** Protein expression of perilipin-1 and PPARγ was measured by immunoblot analysis over the post-differentiation period in WT and SR-BI^-/-^ adipocytes. Tubulin served as a loading control. Representative immunoblots are shown from n=3 independent experiments. **(D)** Secreted adiponectin into the culture medium from WT and SR-BI^-/-^ adipocytes was measured over the post-differentiation period using an ELISA kit. Data are presented as mean ± SD (n=3-4 independent experiments, two-way ANOVA, Tukey’s post hoc [**p≤0.01, ***p≤0.001, ****p≤0.0001]).

Adipose tissue is known to function as an endocrine organ and secretes adipokines (e.g., adiponectin) for signaling and maintaining whole-body metabolic homeostasis. We next tested whether WT and SR-BI^-/-^ adipocytes differentiated from EMSCs had the capability to secrete adiponectin into the culture medium. Both genotypes showed a significant increase in secretion of adiponectin into the culture medium at days 7 and 9 post-differentiation compared to undifferentiated cells at day 0 (**Figure 2D**). Collectively, data presented in Figures 1 and 2 indicate that EMSCs differentiated into adipocytes from WT and SR-BI^-/-^ mice exhibit adipocyte-like characteristics, and therefore can be used as a model system to study adipocyte biology *in vitro*.

### SR-BI trafficking is sensitive to insulin in WT EMSC-derived adipocytes

Increased cell surface expression of SR-BI in response to insulin has been reported in 3T3-L1 adipocytes^21,24^, primary adipocytes^21^, and hepatocytes^27^. To measure cell surface expression of SR-BI in response to insulin in our model system, we treated WT and SR-BI^-/-^ EMSC-derived adipocytes with 100 nM insulin for 30 and 60 min followed by incubation with NHS-linked biotin for subsequent immunoprecipitation of cell surface proteins. As expected, WT EMSC-derived adipocytes that were treated with insulin showed an increase in biotinylated cell surface expression of SR-BI over a one-hour period compared to unstimulated cells (**Figures 3A** and **3B**). SR-BI^-/-^ adipocytes were used as a control for antibody specificity. Further, these data indicate that EMSCs isolated from genetically modified mice can be differentiated into adipocytes and do not require additional knockdown strategies, as we observed no protein expression of SR-BI in SR-BI^-/-^ adipocytes (**Figure 3**).

**Figure 3.**
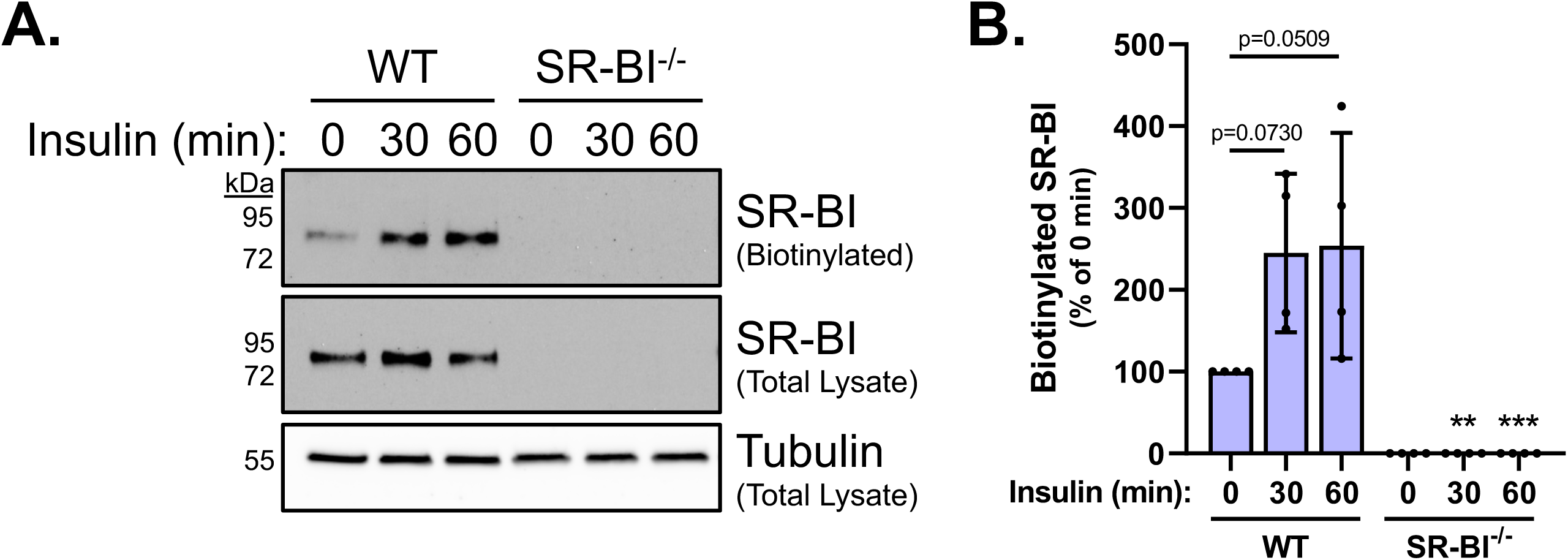
SR-BI trafficking is responsive to insulin in EMSC-derived adipocytes. **(A)** WT and SR-BI^-/-^ adipocytes were serum-starved for 3-4 h in DMEM/0.5% BSA followed by a 30 and 60 min incubation with insulin (100 nM final concentration) at 37°C. Cell surface proteins were labeled with EZ-Link sulfo-NHS-LC-biotin for 1 h at 4°C and immunoprecipitated using streptavidin beads. Biotinylated SR-BI expression (in 100 µL total lysate, mean ± SD of the actual protein concentration = 1.42 ± 0.23 µg/µL for 24 total samples) and total SR-BI expression were measured by immunoblot using an antibody directed against SR-BI. Tubulin served as a loading control for total lysate. Representative immunoblots were selected. **(B)** Densitometry analysis of biotinylated SR-BI over a one-hour period was normalized as a percentage of SR-BI’s expression at 0 min (0 min = 100%). Data are presented as mean ± SD (n=4 independent experiments, two-way ANOVA, Tukey’s post hoc [**p≤0.01 vs. WT 30 min, ***p≤0.001 vs. WT 60 min]).

### SR-BI is required for efficient glucose uptake in adipocytes

With a validated adipocyte model system in hand, we were able to test our hypothesis that SR-BI is required for insulin-stimulated glucose uptake. To begin these studies, we deprived WT and SR-BI^-/-^ adipocytes of glucose using Krebs-Ringer phosphate HEPES (KRPH) buffer for 30 min followed by incubation with 0.5 µCi/well [^3^H]-2-deoxy-D-glucose (2DG) and 50 µM “cold” 2DG with or without 100 nM insulin for 15 min. As expected, incubating WT adipocytes with insulin stimulated a significant 21% increase in 2DG uptake compared to unstimulated adipocytes (**Figure 4A**). In the absence of SR-BI, adipocytes retained the ability to mediate insulin-stimulated glucose uptake, as shown by a significant 27% increase in glucose uptake compared to their unstimulated controls (**Figure 4A**). Despite the similar responsiveness to insulin to mediate glucose uptake in WT and SR-BI^-/-^ adipocytes, we observed that SR-BI^-/-^ adipocytes had a reduced ability to take up glucose compared to WT under basal (**Figure 4B**) and insulin-stimulated (**Figure 4C**) conditions, as shown by a 17% and 13% decrease, respectively. These data indicate that SR-BI is likely not required for adipocytes to respond to insulin to facilitate glucose uptake, but may be required for efficient glucose uptake.

**Figure 4.**
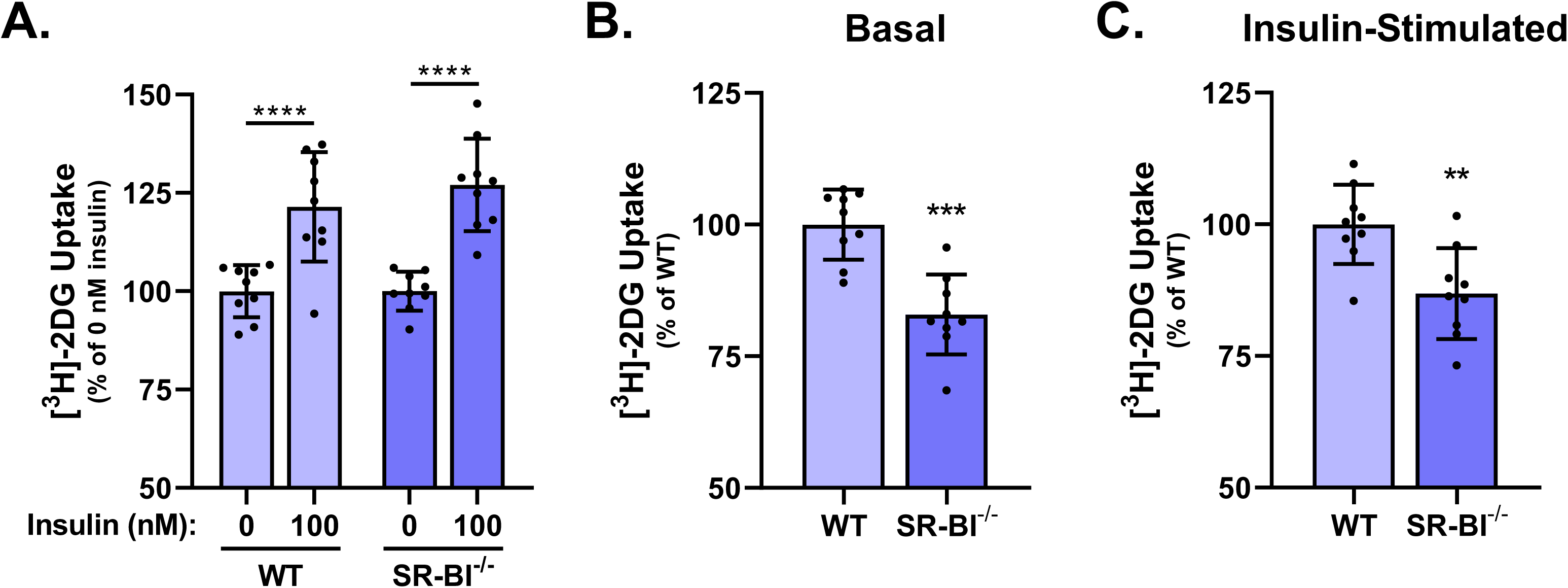
Loss of SR-BI in adipocytes results in inefficient glucose uptake compared to WT adipocytes. **(A)** WT and SR-BI^-/-^ adipocytes were serum-starved for 3-4 h in DMEM/0.5% BSA followed by a 30 min incubation in KRPH buffer. A radioactive solution containing ± insulin (100 nM final concentration), 0.5 µCi/well [^3^H]-2DG, “cold” 2DG (50 µM final concentration) in KRPH buffer was spiked into each well and incubated for 15 min. Cells were lysed in RIPA buffer and radioactivity was measured for each lysate using a scintillation counter. Data from Panel A were reanalyzed to compare **(B)** basal and **(C)** insulin-stimulated glucose uptake compared to WT. DPM values were initially normalized to total protein followed by additional normalization as a percentage of unstimulated controls for each individual genotype (0 nM insulin = 100%) or WT (WT = 100%) as indicated in each panel. Data are presented as mean ± SD (n=3 independent experiments performed in triplicate). Panel A: two-way ANOVA, Tukey’s post hoc (****p≤0.0001), Panels B and C: unpaired Student’s t-test (**p≤0.01, ***p≤0.001).

### Loss of SR-BI in adipocytes results in increased cell surface expression of GLUT4

Translocation of GLUT4 from intracellular storage vesicles to the cell surface in response to insulin is a well-established phenomenon that occurs in adipocytes^28,29^ and muscle^30^. In adipocytes, increased cell surface expression of GLUT4 in response to insulin allows for the uptake and metabolism of excess dietary glucose molecules. The inefficient glucose uptake we observed in SR-BI^-/-^ adipocytes compared to WT adipocytes prompted us to measure the expression of GLUT4 at the cell surface. As expected, when WT adipocytes were treated with insulin for 1 h, cell surface expression of GLUT4 was 35% higher than untreated cells (**Figure 5A**). However, loss of SR-BI resulted in the inability of GLUT4 to translocate to the cell surface of adipocytes in response to insulin compared to their untreated controls (**Figure 5A**). Despite the inability to mediate insulin-stimulated GLUT4 translocation, a re-analysis of the same data demonstrated that SR-BI^-/-^ adipocytes already had significantly higher basal levels of GLUT4 at the cell surface compared to WT adipocytes (**Figure 5B**). These differences in GLUT4 cell surface expression were no longer observed under insulin-stimulated conditions compared to WT adipocytes (**Figure 5C**). We also measured gene expression of *Slc2a4* (GLUT4) and observed that SR-BI^-/-^ adipocytes had upregulated *Slc2a4* gene expression compared to WT adipocytes (**Figure 5D**). These data suggest that SR-BI may be important for regulating the expression and trafficking of GLUT4 in adipocytes.

**Figure 5.**
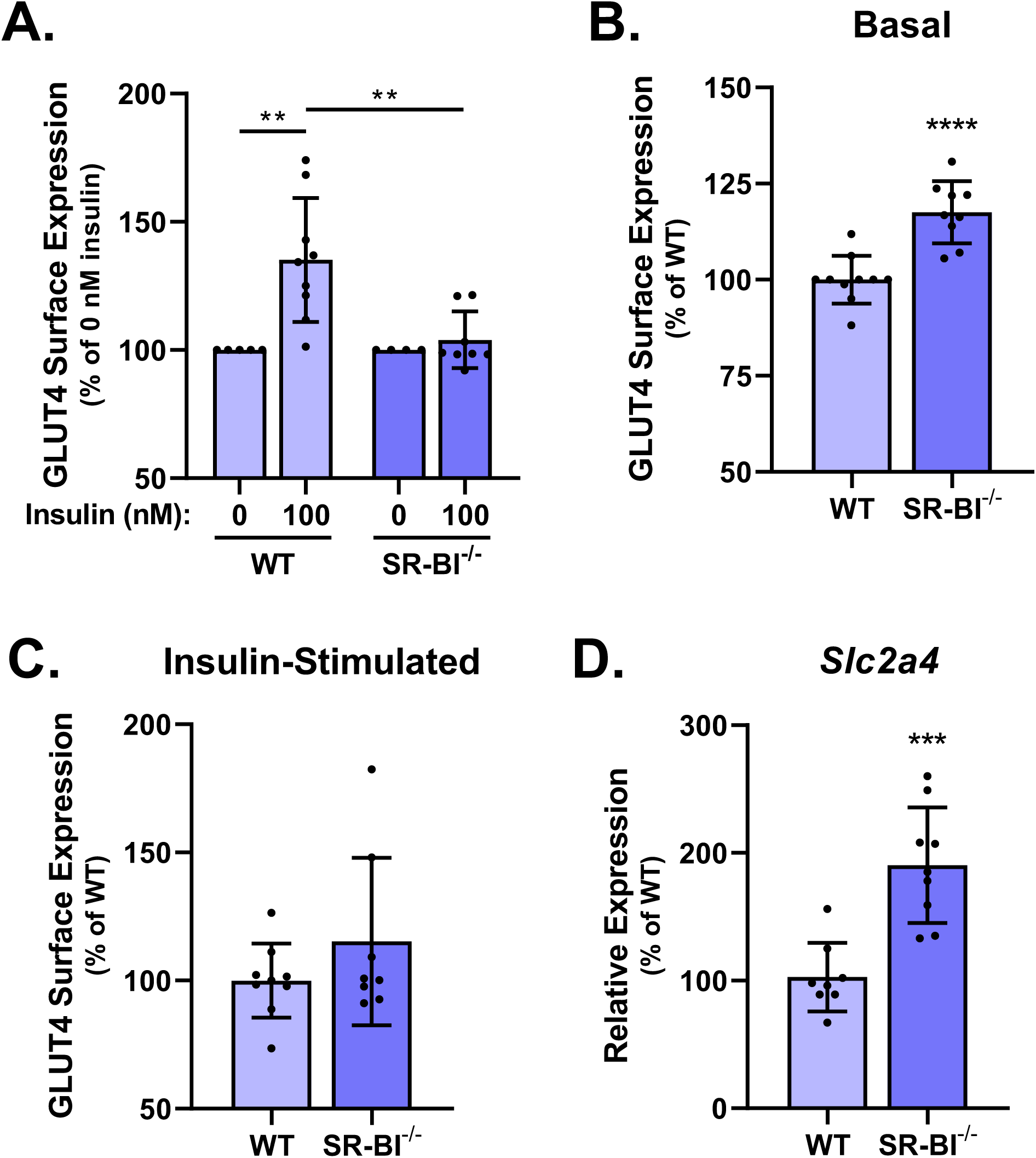
SR-BI may be required for GLUT4 trafficking in adipocytes. **(A)** WT and SR-BI^-/-^ adipocytes were serum-starved for 3-4 h in DMEM/0.5% BSA followed by a 1 h incubation with or without insulin (100 nM final concentration) at 37°C. Cell surface expression of GLUT4 was measured by single-cell suspension FACS analysis using a PE-GLUT4 antibody. Data from Panel A were reanalyzed to measure **(B)** basal and **(C)** insulin-stimulated GLUT4 cell surface expression compared to WT adipocytes. Geometric means from each sample were normalized as a percentage of unstimulated controls for each individual genotype (0 nM insulin = 100%) or WT (WT = 100%) as indicated in each panel. Data are presented as mean ± SD (n=4-5 independent experiments performed in single or duplicate). Panel A: two-way ANOVA, Tukey’s post hoc; Panels B and C: unpaired Student’s t-test (**p≤0.01, ****p≤0.0001). **(D)** Gene expression of *Slc2a4* (GLUT4) was measured in WT and SR-BI^-/-^ adipocytes by the 2^-ΔΔCt^ method using *Rplp0* (60S ribosomal subunit) as a housekeeping gene. Data were then subsequently normalized as a percentage of WT (WT = 100%) and are presented as mean ± SD (n=3 independent experiments performed in duplicate or triplicate, unpaired Student’s t-test [***p≤0.001]).

### SR-BI^-/-^ adipocytes have a lower glycolytic capacity compared to WT adipocytes

Insulin-stimulated uptake of dietary glucose molecules and their subsequent metabolism (i.e., glycolysis) in adipocytes help supply the basic building blocks necessary for the storage of triglycerides or the generation cellular energy in the form of ATP. After observing inefficient glucose uptake in SR-BI^-/-^ adipocytes, we questioned whether the loss of SR-BI impacted the glycolytic capacity of these cells. To measure glycolytic capacity in response to insulin, we performed a Seahorse Glycolytic Rate Assay with an acute injection of insulin into the assay medium (100 nM final concentration) (**Figure 6A**). The Glycolytic Rate Assay accounts for any extracellular acidification (ECAR) that may be coming from oxidative phosphorylation (OXPHOS) by simultaneously measuring oxygen consumption rate (OCR) and ECAR. Subtracting OXPHOS-mediated proton efflux (CO_2_ contributed) from total proton efflux results in the glycolytic proton efflux rate (glycoPER). Compared to WT adipocytes, SR-BI^-/-^ adipocytes displayed a significant reduction in glycoPER before (basal glycolysis) and after (induced glycolysis) the addition of insulin. Further, after addition of rotenone/antimycin A (Rot/Ant A), which inhibits complexes I and III of the electron transport chain, compensatory glycolysis was also significantly reduced (**Figure 6B**). These data indicate that SR-BI may play an important role in regulating glycolytic function in adipocytes, potentially due to the decrease in glucose uptake.

**Figure 6.**
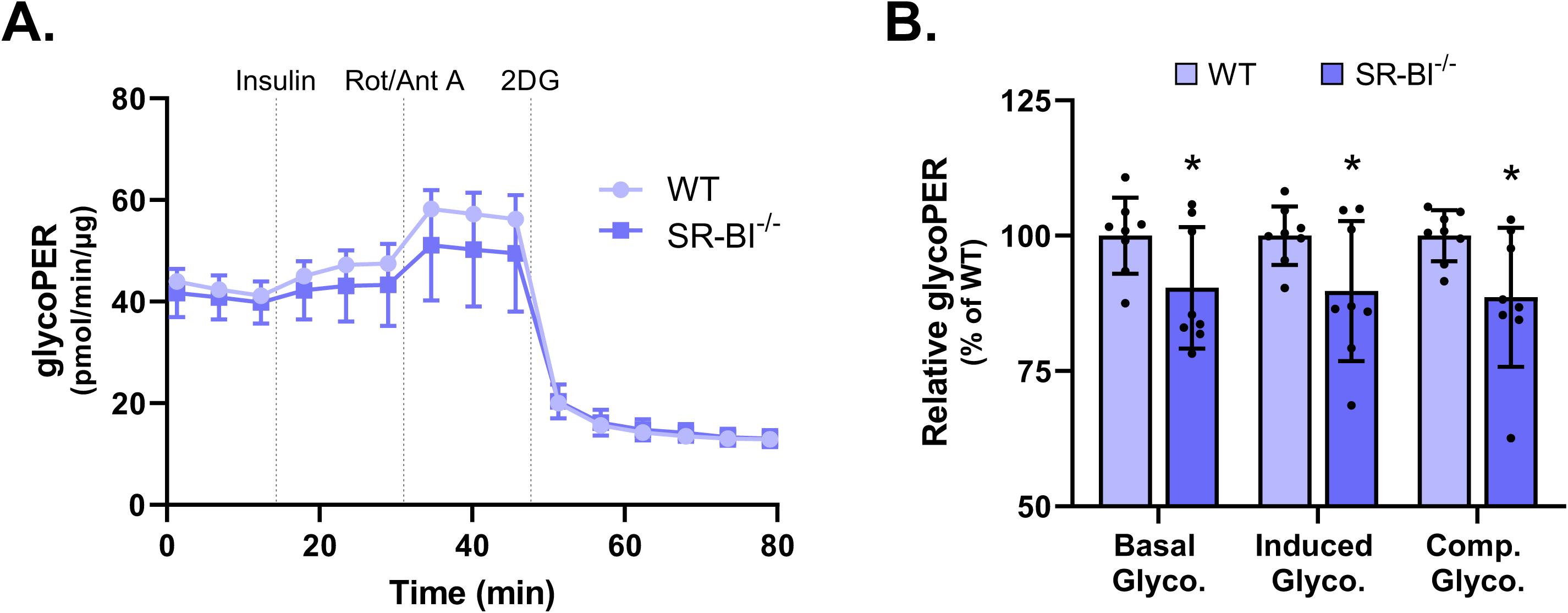
SR-BI deficiency reduces the glycolytic capacity of adipocytes. **(A)** WT and SR-BI^-/-^ adipocytes were serum-starved in DMEM/0.5% BSA for 3-4 h at 37°C followed by a 30-60 min incubation in Seahorse XF DMEM medium at 37°C in a non-CO_2_ incubator. Insulin-stimulated glycolytic proton efflux rate (glycoPER) was measured using the XF Glycolytic Rate Assay performed with an acute insulin injection (100 nM final concentration). Data presented in Panel A are representative of three individual experiments (mean ± SD, normalized to total protein [µg]). **(B)** Measurements from the XF Glycolytic Rate Assay were used to determine basal, induced, and compensatory glycolysis. Data were normalized as a percentage of WT (WT = 100%) and presented as mean ± SD (n=3 independent experiments performed in duplicate or triplicate, unpaired Student’s t-test [*p≤0.05]).

### Insulin-stimulated de novo fatty acid and triglyceride synthesis is increased in SR-BI^-/-^ adipocytes

The intracellular fate of glucose includes the metabolic conversion into lipids, such as fatty acids and triglycerides, to support normal cell function, growth, and homeostasis. One of the intermediates in the glycolytic metabolic pathway is the conversion of glucose to acetyl-coenzyme A (CoA). To measure whether the reduced glucose uptake observed in SR-BI^-/-^ adipocytes impacted *de novo* lipid synthesis, we incubated WT and SR-BI^-/-^ adipocytes with 5 µCi/well [^14^C]- acetic acid for 4 h in the presence or absence of 100 nM insulin and measured the incorporation of radioactivity into fatty acids and triglycerides by thin layer chromatography. SR-BI^-/-^ adipocytes showed significant increases in the incorporation of [^14^C]-acetic acid into fatty acids (**Figure 7A**) and triglycerides (**Figure 7B**), compared to WT adipocytes after insulin stimulation, but not at basal levels. Basal triglyceride synthesis between WT and SR-BI^-/-^ adipocytes was considered significant (p≤0.05) by unpaired Student’s t-test (**Figure 7B**). These data indicate that SR-BI^-/-^ adipocytes may have altered lipid metabolism compared to WT adipocytes by prioritizing fatty acid and triglyceride synthesis after insulin stimulation.

**Figure 7.**
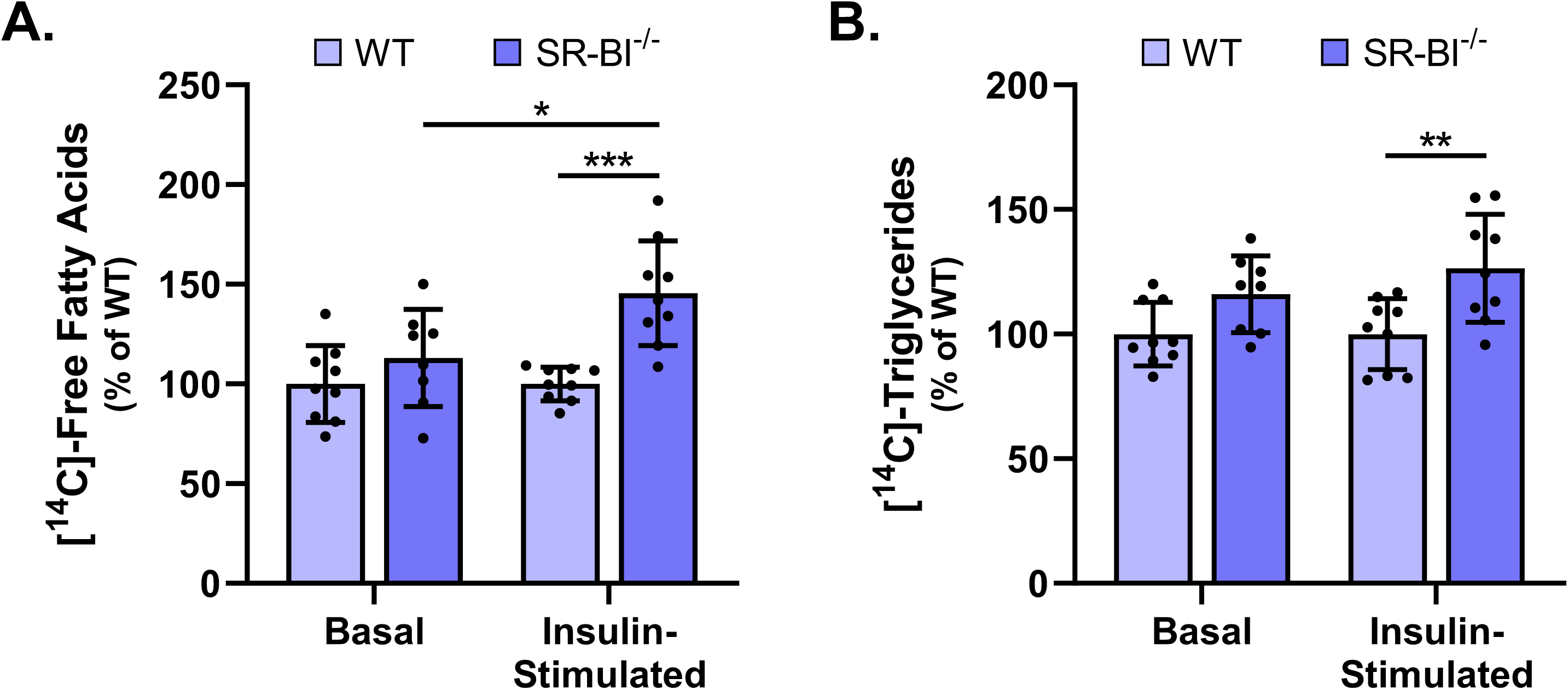
*De novo* lipid synthesis is altered in SR-BI^-/-^ adipocytes compared to WT adipocytes. WT and SR-BI^-/-^ adipocytes were serum starved in DMEM/0.5% BSA for 3-4 h followed by incubation with or without insulin (100 nM final concentration) and 5 µCi/well [^14^C]- acetic acid in DMEM/0.5% BSA containing 50 µM sodium acetate for 4 h. Cellular lipids were extracted with isopropanol and separated on high-performance thin layer chromatography glass silica plates. The incorporated radioactivity in **(A)** free fatty acids and **(B)** triglycerides was measured using a liquid scintillation counter. Data were normalized to total cellular protein and presented as a percentage of WT (WT = 100%, mean ± SD, n=3 independent experiments performed in duplicate or triplicate, two-way ANOVA, Tukey’s post hoc [*p≤0.05, **p≤0.01, ***p≤0.001]).

### SR-BI^-/-^ adipocytes have differential regulation of lipid metabolism genes compared to WT adipocytes

We next sought to determine whether genes related to lipid metabolism (i.e., glucose uptake, fatty acid synthesis, triglyceride synthesis, and β-oxidation) were differently regulated in SR-BI^-/-^ adipocytes compared to WT adipocytes. At basal levels, SR-BI^-/-^ adipocytes showed significant upregulation of *Slc2a4* (GLUT4), *Fasn* (fatty acid synthase [FASN]), and *Dgat1* (diacylglycerol O-acyltransferase 1 [DGAT1]) compared to WT adipocytes (**Figure 8A**). After insulin stimulation, gene expression of *Insr* (insulin receptor [INSR]), *Slc2a4*, *Fasn*, and *Acadl* (long-chain acyl-CoA dehydrogenase [LCAD]) was significantly upregulated in SR-BI^-/-^ adipocytes compared to WT adipocytes (**Figure 8B**). The increase in *Fasn* and *Dgat1* gene expression observed in SR-BI^-/-^ adipocytes may account for the increased incorporation of radioactivity into fatty acids (**Figure 7A**) and triglycerides (**Figure 7B**) compared to WT adipocytes. All other genes measured in basal or insulin-stimulated SR-BI^-/-^ adipocytes showed no statistical differences in expression compared to WT adipocytes.

**Figure 8.**
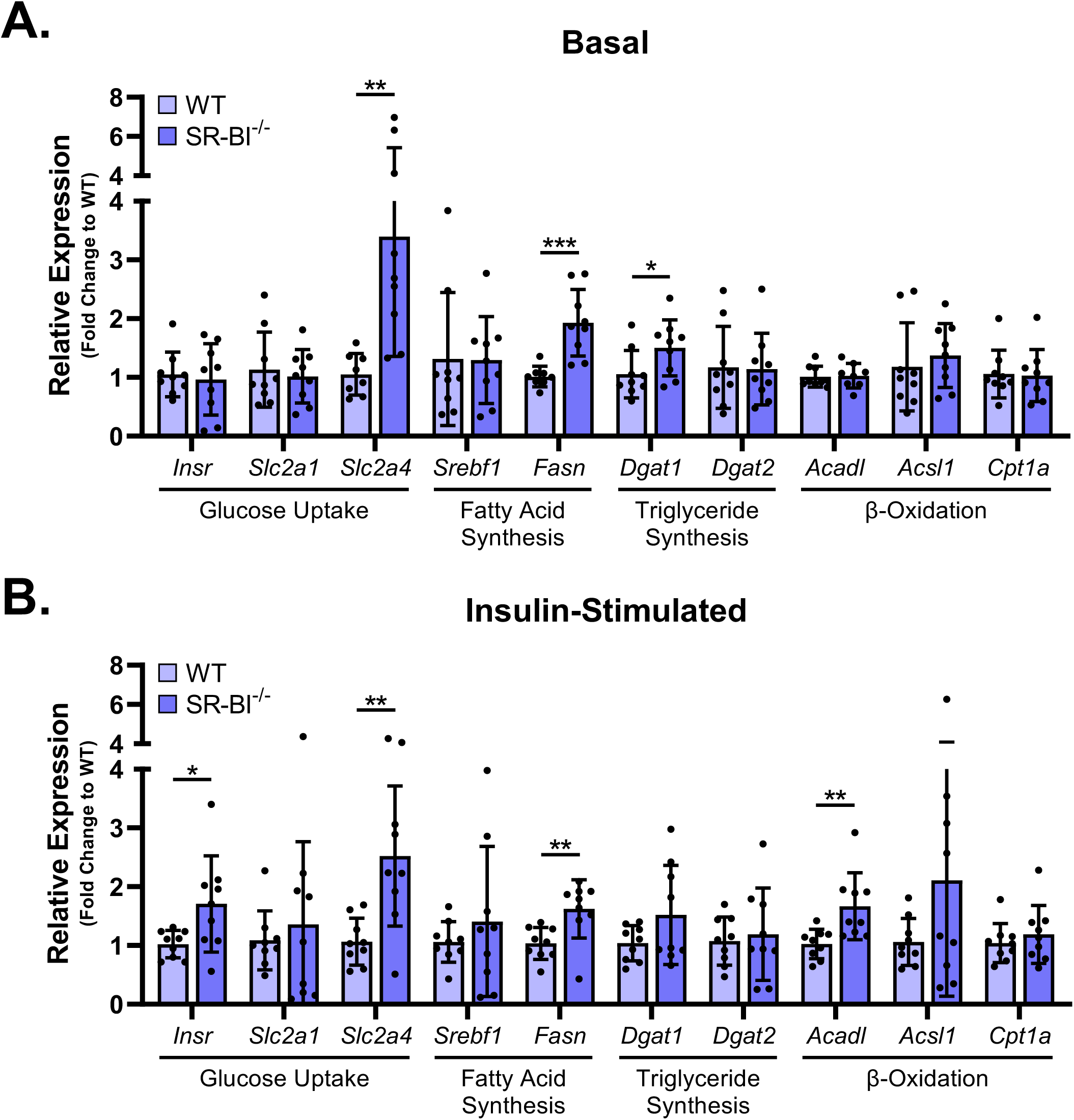
SR-BI^-/-^ adipocytes have dysregulated expression of lipid metabolism genes. WT and SR-BI^-/-^ adipocytes were deprived of insulin for 3-4 h in complete medium. Following insulin deprivation, adipocytes were incubated in the **(A)** absence or **(B)** presence of insulin (100 nM final concentration) for 2 h at 37°C. Expression of genes related to glucose uptake, fatty acid synthesis, triglyceride synthesis, and β-oxidation were analyzed by the 2^-ΔΔCt^ method using *Rplp0* (60S ribosomal subunit) as a housekeeping gene. Data were then subsequently normalized as a fold change to WT (WT = 1) and presented as mean ± SD (n=3 independent experiments performed in duplicate or triplicate, unpaired Student’s t-test [*p≤0.05, **p≤0.01, ***p≤0.001]).

## Discussion

The requirement of SR-BI to mediate glucose uptake by its native ligands, apolipoprotein A-1 (apoA-1) in skeletal muscle^35^ in an insulin-dependent and -independent manner^31^, as well as by HDL in adipocytes^23^ has been reported. The latter study did not test whether loss of SR-BI in adipocytes elicited a change in glucose uptake in the presence of insulin. In this study, we designed experiments to test the hypothesis that SR-BI was required for insulin-stimulated glucose uptake in adipocytes. We demonstrated the usefulness of EMSCs isolated from SR-BI^-/-^ mice to study adipocyte biology *in vitro* as they can be differentiated into adipocytes (Figures 1, 2, and Supplemental Figure 1) and maintain their knockout phenotype (Figure 3). To test our hypothesis, we chose an *in vitro* model system that utilizes EMSCs isolated from WT and SR-BI^-/-^ mice that can be differentiated into adipocytes. EMSCs have proven to be advantageous to other *in vitro* adipocyte model systems because ear tissue collection is a survival procedure; they do not require additional knockdown strategies to study the effect of specific proteins; they are easy to isolate, culture, and differentiate; and simultaneous isolations can occur across various genotypes. Additionally, EMSCs have also been shown to have a greater adipogenic potential compared to murine preadipocytes isolated from the adipocyte stromal vascular fraction^32^.

Using our differentiated EMSC model, we showed that in the absence or presence of insulin, glucose uptake is inefficient in SR-BI^-/-^ adipocytes (Figures 4B and 4C) compared to WT adipocytes, indicating a novel link between adipocyte SR-BI and glucose uptake. To our knowledge, we are the first to demonstrate that SR-BI deficiency in adipocytes results in a reduced glycolytic capacity, as shown by our glycoPER data (Figure 6), most likely due to the decrease in glucose uptake. Further, *de novo* lipogenesis experiments suggested that SR-BI^-/-^ adipocytes may generate more fatty acids and triglycerides than WT adipocytes (Figure 7A), which is supported by upregulated gene expression of *Fasn* and *Dgat1* in SR-BI^-/-^ adipocytes (Figures 8A and 8B). Altogether, the data presented in this manuscript partially support our hypothesis and suggest that SR-BI is required for efficient glucose uptake, regardless of insulin, and maintaining metabolic homeostasis in adipocytes.

Surprisingly, despite decreased levels of glucose uptake, we observed increased gene expression of the insulin-responsive glucose transporter, *Slc2a4* (GLUT4), in SR-BI^-/-^ adipocytes (Figures 5D, 8A, and 8B). This observation has been noted before by Hoekstra et al.^33^ who suggested that increased GLUT4 gene expression would correlate with increased glucose uptake, although the latter was not directly measured in their study. Glucose tolerance testing in SR-BI^-/-^ mice on a high-fat diet (HFD) showed that male and female mice could clear glucose more efficiently than WT mice. This data coincided with an increase in total and gonadal fat mass in the SR-BI^-/-^ mice when fed a HFD. Together, the increased rate of glucose clearance with SR-BI deficiency may be due to increased glucose uptake in adipose tissue, thereby predisposing SR-BI^-/-^ mice to obesity. On the contrary, another study found that loss of SR-BI in mice did not lead to increased weight gain compared to HFD-fed WT mice, although it was observed that SR-BI^-/-^ mice had hypertrophied adipocytes^34^. It is possible that glucose clearance in SR-BI^-/-^ mice is mediated by skeletal muscle, as skeletal muscle was found to be responsible for the majority of insulin-stimulated whole-body glucose clearance^35^, although this does appear to be dependent on species, administration method, concentration, fed state, and obesogenic state^36,37^. The discrepancy between glucose uptake *in vivo* versus *in vitro* could be explained by the fact that mice have global SR-BI deficiency and the effects of this genetic background may impact glucose uptake into other organs.

The lack of correlation between GLUT4 cell surface expression and inefficient glucose uptake in SR-BI^-/-^ adipocytes is perplexing. It has been previously reported that after the removal of insulin, 50% of GLUT4 in adipocytes gets rapidly internalized and is nearly absent from the plasma membrane within 20 min^38^. In the absence of insulin, we observed increased cell surface expression of GLUT4 in SR-BI^-/-^ adipocytes compared to WT adipocytes (Figure 5B). In fact, GLUT4 recycling may be defective in SR-BI^-/-^ adipocytes since we observed no change in GLUT4 cell surface expression after insulin stimulation (Figure 5A). Cholesterol and fatty acid content within the plasma membrane mediates membrane fluidity, as well as membrane organization through the formation of lipid raft microdomains. Given SR-BI’s role in cholesterol transport^20,21,39,40^ and fatty acid uptake^22^ in adipocytes, it is possible that GLUT4 recycling in SR-BI^-/-^ adipocytes may be dysfunctional due to disrupted lipid transport and lipid raft formation, which has been shown to be important for GLUT4 trafficking^41^. A study performed by Gustavsson, et al.^42^ demonstrated that insulin stimulated a rapid increase of GLUT4 to the cell surface, but glucose uptake did not occur until GLUT4 was present in caveolae-rich microdomains of the plasma membrane. If lipid raft formation is reduced in SR-BI^-/-^ adipocytes, GLUT4 may be unable to transport glucose efficiently similar to what we observed in our glucose uptake studies (Figure 4). Additionally, studies from Shigematsu, et al.^43^ and Ros-Baró, et al.^41^ demonstrated that the rate at which adipocyte GLUT4 endocytosed in the absence of insulin was dramatically reduced with altered lipid raft formation and caveolin-1 assembly. It was also observed that increased basal accumulation of GLUT4 at the cell surface was due to defective GLUT4 endocytosis^43^, which may account for the increase of GLUT4 cell surface expression we observed in SR-BI^-/-^ adipocytes under basal conditions compared to WT adipocytes (Figure 5B). Further studies are required to establish the direct role that SR-BI may play in GLUT4 recycling, plasma membrane localization, and function.

We observed altered expression of lipid metabolism genes in the absence (Figure 8A) and presence (Figure 8B) of insulin in SR-BI^-/-^ adipocytes compared to WT adipocytes. Specifically, we demonstrated upregulation of *Fasn* and *Dgat1* gene expression in SR-BI^-/-^ adipocytes compared to WT adipocytes, which may account for the increased incorporation of [^14^C]-acetic acid into fatty acids (Figure 7A), and triglycerides (Figure 7B), respectively. Wang, et al.^22^ showed that SR-BI is required for fatty acid uptake in adipocytes. The observed increase in *Fasn* gene expression and fatty acid synthesis in SR-BI^-/-^ adipocytes is potentially a compensatory mechanism to retain the ability to synthesize triglycerides for excess fat storage or for ATP production through OXPHOS.

Within the last several years, a protein partnership between SR-BI and an extracellular matrix protein called procollagen C-endopeptidase enhancer 2 (PCPE2) was established in hepatocytes^40^ and adipocytes^39,44^. On an atherosclerotic low-density lipoprotein receptor knockout (LDLR^-/-^) background, hepatic^40^ and adipocyte^44^ SR-BI had a reduced ability to facilitate HDL binding and cholesteryl ester uptake from HDL into cells in the absence of PCPE2. While the direct mechanisms by which PCPE2 may facilitates SR-BI’s ability to transport cholesterol continue to be investigated, it is possible that SR-BI and PCPE2 may partner to facilitate other metabolic processes in adipocytes, including glucose uptake and/or downstream metabolic processes and warrant further investigation.

In **Figure 9**, we have highlighted key findings and present a model for what may be occurring in SR-BI^-/-^ adipocytes. Insulin-stimulated glucose uptake in adipocytes is primarily facilitated by GLUT4 at the cell surface. It has been previously demonstrated that efficient glucose uptake and GLUT4 endocytosis are reliant on caveolae-rich microdomains in the plasma membrane^41–43^, which are highly abundant in cholesterol. Considering SR-BI’s role in mediating cholesterol transport in adipocytes^20,21^, it is possible that, in the absence of SR-BI, the reduced ability to transport HDL-CE into adipocytes results in a delipidated membrane. As such, GLUT4 is retained at the cell surface in the absence of insulin but has a reduced ability to transport glucose efficiently. Because of the reduced glucose uptake in SR-BI^-/-^ adipocytes (Figure 4), the ability for these cells to perform glycolysis decreases (Figure 6), thus impacting downstream metabolic processes (Figures 7 and 8). SR-BI has been associated with facilitating fatty acid uptake^22^. As such, the potential decrease in fatty acid uptake with SR-BI deficiency may increase the gene expression of *Fasn* (FASN) (Figure 8) to synthesize new fatty acids (Figure 7). The increase in fatty acid synthesis could be used to fuel OXPHOS or lipid storage as triglycerides, which aligns well with the observed increase in *Dgat1* (DGAT1) gene expression (Figure 8) and increased triglyceride synthesis (Figure 7) in SR-BI^-/-^ adipocytes. Lastly, it is possible that the increase in *Acadl* (LCAD) gene expression (Figure 8), an enzyme that catalyzes β-oxidation of long-chain fatty acids^45^, could be fueling OXPHOS. Whether loss of SR-BI induces a metabolic switch from glycolysis to OXPHOS in adipocytes requires additional investigation. These hypotheses are thought-provoking and set the stage for SR-BI being a potential therapeutic target for obesity and its related comorbidities.

**Figure 9.**
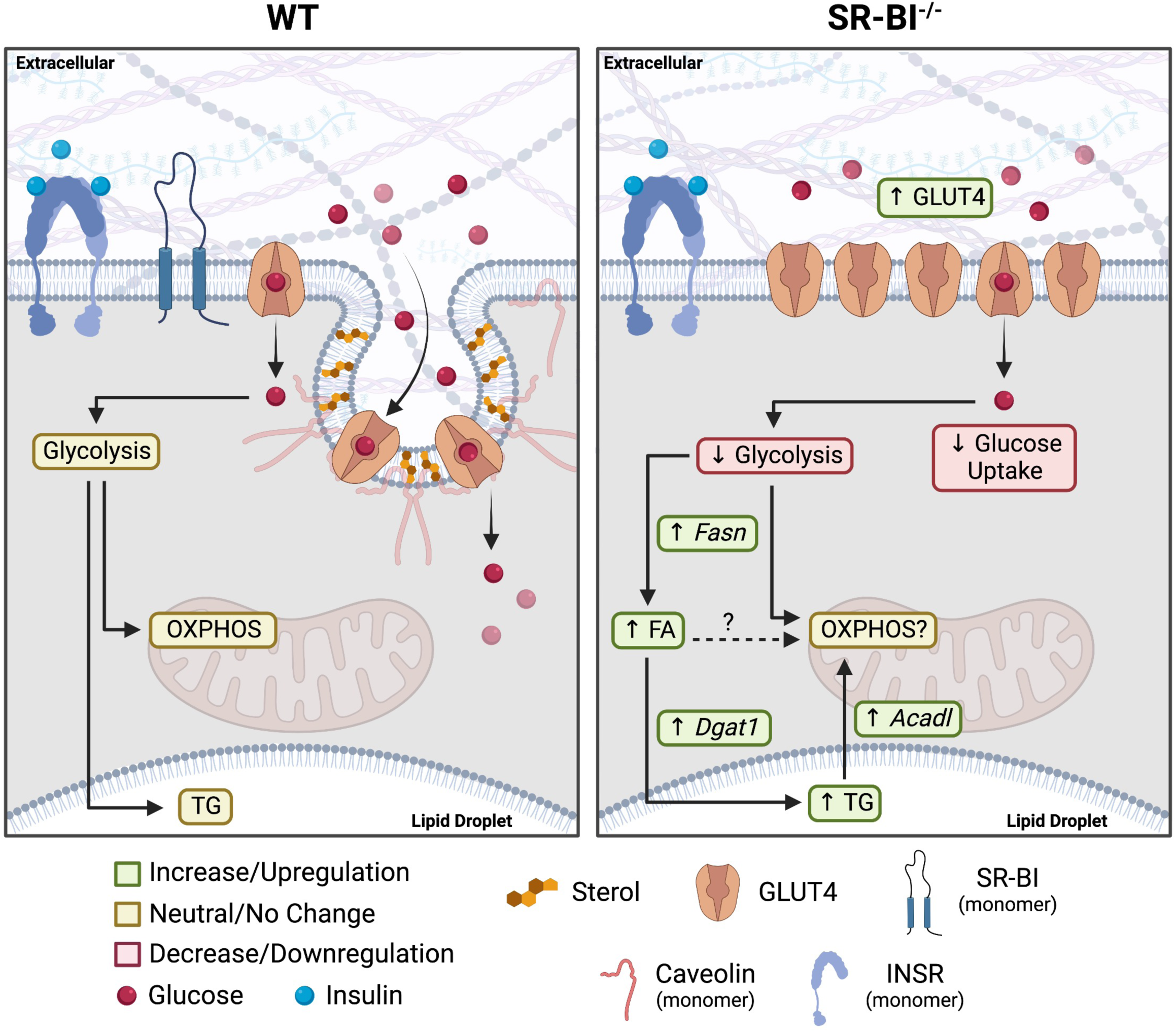
Proposed model describing glucose and lipid metabolism in WT and SR-BI^-/-^ adipocytes. (**Left**) Insulin-stimulated glucose uptake is facilitated by GLUT4 that is localized to caveolae- and cholesterol-rich lipid raft microdomains within the plasma membrane. Additionally, these lipid raft microdomains have been shown to be important for GLUT4 internalization in the absence of insulin. The intracellular fates of glucose in adipocytes include energy supply through glycolysis and/or OXPHOS and lipid storage in the form of triglycerides. (**Right**) In the absence of SR-BI, cholesterol transport in adipocytes is decreased, thus creating a delipidated membrane. Despite GLUT4 being at the cell surface, decreased lipid raft formation is unable to support efficient glucose uptake under basal and insulin-stimulated conditions, ultimately leading to a reduced glycolytic capacity. Lastly, SR-BI-deficient adipocytes upregulate *Fasn* and *Dgat1* gene expression to synthesize fatty acids and triglycerides for lipid storage, possibly as a compensatory mechanism for reduced fatty acid and glucose uptake. INSR: insulin receptor. Illustration not to scale. Created with BioRender.

Our lab, and others, have extensively characterized cholesterol transport functions of genetic human variants of SR-BI^46,47^ and cardiovascular risk^48–50^. Surprisingly, SR-BI single nucleotide polymorphisms and their association with metabolic diseases are understudied. With growing evidence that SR-BI is important for adipocyte function, measuring the impact these genetic variants of SR-BI have on adipocyte function could potentially shed light on the risk of obesity, metabolic syndrome, and insulin sensitivity.

## Materials and Methods

### Materials

Probucol, dexamethasone, 3-isobutyl-1-methylxanthine (IBMX), human insulin, and bovine insulin were purchased from Sigma-Aldrich (St. Louis, MO, USA). Dulbecco’s Modified Eagle Medium/Nutrient Mixture F-12 (DMEM/F12), collagenase IV, human insulin, TRIzol, DNase I, high-capacity complementary DNA (cDNA) reverse transcriptase kit, SYBR green, EZ-Link sulfo-NHS-LC-Biotin, and high-capacity streptavidin beads were purchased from Thermo Fisher Scientific (Waltham, MA, USA). Rosiglitazone was purchased from Cayman Chemical (Ann Arbor, MI, USA). 2-[1,2-^3^H(N)]-deoxy-D-glucose and [1,2-^14^C]-acetic acid were purchased from PerkinElmer (Waltham, MA, USA). PPARγ, perilipin-1, and ɑ-tubulin antibodies were purchased from Cell Signaling Technology (Danvers, MA, USA). SR-BI antibody and mouse adiponectin ELISA kit was purchased from Abcam (Cambridge, UK, England). PE-GLUT4 antibody was purchased from Santa Cruz Biotechnology (Dallas, TX, USA). 2-deoxy-D-glucose (2DG) was purchased from Alfa Aesar (Haverhill, MA, USA). Primers for quantitative PCR were designed and purchased from IDT Technologies (Coralville, IA, USA). All other reagents were of analytical grade.

### Animal Care

Experimental procedures performed in mice conformed to the Guide for the Care and Use of Laboratory Animals (NIH publication, 8^th^ Edition, 2011) and were approved by the Institutional Animal Care and Use Committee of the Medical College of Wisconsin. WT and SR-BI^-/-^ mice, both C57BL6/J strain, were housed and maintained under a normal light-dark cycle in a pathogen-free barrier facility in accordance with federal and institutional guidelines. SR-BI^-/-^ breeders were fed a 0.5% probucol diet (Envigo, Indianapolis, IN, USA) to restore female fertility that is lost with SR-BI deficiency^51^. At the time of EMSC isolation, mice were euthanized by CO_2_ inhalation followed by cervical dislocation. WT mice were originally purchased from Jackson Laboratories (Bar Harbor, ME, USA) and SR-BI^-/-^ ^52^ mice were received as a gift from Dr. Monty Krieger (MIT, Cambridge, MA, USA).

### Isolation and Differentiation of EMSCs

EMSCs were isolated from WT and SR-BI^-/-^ mice as previously described^32,53^ with minor modifications. Briefly, the outer ears of male or female mice between 8-12 weeks of age were excised, minced, and enzymatically digested with 1.5 mg/mL collagenase IV (245 units/mg) at 37°C in a shaking water bath for 1 h at 200 rpm. EMSCs were cultured at 37°C/5% CO_2_ in growth medium (DMEM/F12, 15% fetal bovine serum (FBS), 1% penicillin-streptomycin (P/S)) supplemented with 10-100 ng/mL recombinant murine fibroblast growth factor (Peprotech, Rocky Hill, NJ, USA). At passage 3, cells were plated for experiments and allowed to reach maximum confluency and incubated for an additional 48 h in growth medium. At day 0 (4 days total after initial plating), cells were cultured in differentiation medium (growth medium, 5 µg/mL insulin, 3 µM rosiglitazone, 1 µM dexamethasone, 500 µM IBMX). After day 2 of post-differentiation, cells were fed every other day with maintenance medium (growth medium, 5 µg/mL insulin, 3 µM rosiglitazone) until day 9 post-differentiation. Experiments described in this study were performed in adipocytes that were between day 7 to day 9 post-differentiation and ranging from passage 3 to passage 5, unless otherwise stated.

### Oil Red O Staining, Imaging, and Quantification

Cells at various days post-differentiation were fixed with 4% paraformaldehyde for 15 min at room temperature (RT) followed by staining with ORO for 5 min. Stained cells were imaged on an Olympus LS CKX53 microscope (Waltham, MA, USA) at 20X magnification. Three different fields of view were acquired for each sample and representative images were selected. For quantification, ORO stain was extracted using isopropanol for 5 min at RT and absorbance was measured at 500 nm on a plate reader. A tissue culture plate without cells was stained with ORO and extracted as described above to serve as a “blank” measurement.

### Adiponectin Secretion by ELISA

Cultured medium was collected 18-24 h post-medium change from cells at days 0, 7, and 9 post-differentiation and cleared via centrifugation at 3,000 *x g* for 10 min at 4°C. Biological replicates (n=3) from individual experiments were combined before analyzing the amount of secreted adiponectin into the culture medium using a mouse adiponectin ELISA kit following manufacturer’s instructions.

### Cell Lysis

Total cellular proteins were harvested from undifferentiated EMSCs (day 0) or adipocytes at various days post-differentiation by incubating with radioimmunoprecipitation assay (RIPA) buffer (1% NP-40 buffer, 10 mM Tris [pH 8.0], 150 mM NaCl, 0.5% sodium deoxycholate, 0.1% SDS) containing protease inhibitors (1 µg/mL leupeptin/aprotinin/pepstatin, 20 µg/mL PMSF) for 10 min on ice. Lysates were subjected to syringe sheering with a 23G needle or pipette trituration followed by centrifugation at 21,130 x *g* for 10 min at 4°C. Protein concentrations were determined via the Lowry method^54^.

### Immunoblot Analysis

Cellular lysates (20 µg protein) from adipocytes at various days post-differentiation were separated via SDS-PAGE and wet-transferred to nitrocellulose membrane. Protein expression was measured using antibodies directed towards PPARγ (1:1,000), perilipin-1 (1:1,000), SR-BI (1:1,000), and tubulin (1:5,000). Bands were visualized using the appropriate secondary antibodies and ECL substrate on a BioRad ChemiDoc system (Hercules, CA, USA) or x-ray film. Band intensities were quantified using NIH FIJI/ImageJ software. Protein expression quantification was determined using tubulin as a loading control.

### RNA Extraction and Reverse Transcriptase PCR

RNA from adipocytes at various days post-differentiation was extracted using the TRIzol phenol-chloroform method according to manufacturer’s instructions with some modifications. Briefly, cells were lysed with TRIzol for 5 min followed by two consecutive 3 min incubations with chloroform at RT. Each wash was followed by centrifugation at 12,000 x *g* for 10 min at 4°C. The aqueous layer was transferred to a new 1.5 mL tube and incubated with RT isopropanol for 10 min. Following centrifugation at 12,000 x *g* for 10 min at 4°C, the isopropanol was removed, and the RNA pellet was washed with ice-cold 75% ethanol twice followed by centrifugation at 7,500 x *g* for 5 min at 4°C. RNA pellets were air-dried for 15 min, resuspended in RNase/DNase-free water, and incubated in a 55°C water bath for 10 min. RNA concentrations, A280/260, and A260/230 ratios were read using an ND-1000 Spectrophotometer (Nanodrop, Wilmington, DE, USA). RNA (0.7 – 1 µg) was treated with DNase I and synthesized into cDNA using a high-capacity cDNA reverse transcriptase kit in an Eppendorf Mastercycler EP Gradient Thermal Cycler.

### Quantitative Real-Time PCR Analyses

Quantification of cDNA products was determined using SYBR Green and a 384-well BioRad PCR Detection System. Relative gene expression was determined by the 2^-ΔΔCt^ method with a 40-cycle threshold limit. The 60S ribosomal subunit (*Rplp0*) was used as the housekeeping gene for all quantitative analyses. Melt curves and no reverse transcriptase controls were used for PCR product specificity of each primer set and genomic DNA contamination, respectively. Mouse forward (For) and reverse (Rev) primer sequences (5’ → 3’) used for quantitative PCR include: *Acadl* – For: GGTGGAAAACGGAATGAAAGG, Rev: GGCAATCGGACATCTTCAAAG; *Acsl1* – For: CCTGTGGGATAAACTCATCTTCC, Rev: GTCCGTAGCCTTCATAGAACTG; *Cebpa* – For: GCAAGCCAGGACTAGGAGAT, Rev: AATACTAGTACTGCCGGGCC; *Cpt1a* – For: GATCTACAATTCCCCTCTGCTC, Rev: AGCCAGACCTTGAAGTAACG; *Dgat1* – For: AACTCAACTTTCCTCGGTCC, Rev: GGCTTCATGGAGTTCTGGATAG; *Dgat2* – For: CGAGACTACTTTCCCATCCAG, Rev: AAGTTACAGAAGGCACCCAG; *Fasn* – For: CCCCTCTGTTAATTGGCTCC; Rev: TTGTGGAAGTGCAGGTTAGG; *Insr* – For: CAAGATTCCCCAGATGAGAGG; Rev: AGATTTCATGAGTCACAGGGC; *Pparg2* – For: TGTTATGGGTGAAACTCTGGG, Rev: AGAGCTGATTCCGAAGTTGG; *Pref1* – For: TGTCAATGGAGTCTGCAAGG, Rev: ATTCGTACTGGCCTTTCTCC; *Rplp0* – For: TGACATCGTCTTTAAACCCCG, Rev: TGTCTGCTCCCACAATGAAG; *Slc2a1* – For: GATTGGTTCCTTCTCTGTCGG, Rev: CCCAGGATCAGCATCTCAAAG; *Slc2a4* – For: CATTCCCTGGTTCATTGTGG, Rev: GAAGACGTAAGGACCCATAGC; *Srebf1* – For: GAACCTGACCCTACGAAGTG, Rev: TTTCATGCCCTCCATAGACAC.

### Glucose Uptake

Adipocytes were serum-starved in DMEM/0.5% BSA for 3-4 h followed by a 30 min incubation in KRPH buffer (5 mM Na_2_HPO_4_, 20 mM HEPES, pH 7.4, 1 mM MgSO_4_, 1 mM CaCl_2_, 136 mM NaCl, 4.7 mM KCl, and 1% BSA). After 30 min, a radioactive solution consisting of 0.5 µCi/well of 2-[1,2-^3^H(N)]-deoxy-D-glucose, “cold” 2DG (50 µM final concentration), and with or without insulin (100 nM final concentration) in KRPH buffer was spiked into each well and incubated for 15 min. The radioactive solution was aspirated and each well was washed with cold PBS. Cells were lysed with RIPA buffer for 10 min. The amount of radioactivity in the cell lysate was measured using a liquid scintillation counter (Beckman Coulter, Brea, CA, USA) and protein concentration was determined via the Lowry method^54^. Data were normalized to total protein and as indicated in the figure legend.

### SR-BI Cell Surface Biotinylation

WT and SR-BI^-/-^ adipocytes were serum-starved for 3-4 h in DMEM/0.5% BSA followed by incubation with 100 nM insulin for 30 or 60 min. Cells were then incubated with 1 mg/mL nonmembrane-permeable EZ-Link sulfo-NHS-LC-Biotin in PBS for 1 h at 4°C in the dark. Lysates were collected using 1% NP-40 containing protease inhibitors. Equal volume of biotinylated proteins (100 µL cell lysate, mean ± SD of the actual protein concentration = 1.42 ± 0.23 µg/µL for 24 total samples) were immunoprecipitated using high-capacity streptavidin beads, eluted in 2X Laemmli buffer (4% SDS, 20% glycerol, 120 mM Tris-HCl [pH 6.8], 5% β-mercaptoethanol [BME], bromophenol blue), and separated via SDS-PAGE. Cell surface SR-BI (biotinylated), total SR-BI, and tubulin (loading control for total lysate) were detected by immunoblot.

### GLUT4 Cell Surface Expression

Adipocytes (days 7-10 post-differentiation) were serum-starved for 3-4 h in DMEM/0.5% BSA followed by a 1 h incubation with or without 100 nM insulin. Following incubation, adipocytes were gently scraped from the plate, resuspended in PE-GLUT4 primary antibody (1:1,000) prepared in fluorescence-activated cell sorting (FACS) buffer (PBS, 5% FBS, 0.1% sodium azide), and incubated at RT in the dark for 15 min. The cell suspension was centrifuged at 1,000 *x g* for 1 min, resuspended in FACS buffer, and analyzed on a BD LSRII flow cytometer (Franklin Lakes, NJ, USA). Geometric means from each sample were normalized as indicated in each panel.

### Seahorse Assays

WT and SR-BI^-/-^ EMSCs were plated in Agilent Seahorse XF96 V3 PS Cell Culture Microplates (Santa Clara, CA, USA) at a density of 3,000 cells/well. Four days after initial plating, EMSCs were differentiated as previously described. Adipocytes were serum-starved for 3-4 h in DMEM/0.5% BSA followed by incubation in Agilent Seahorse XF assay medium (DMEM, 1 mM pyruvate, 2 mM glutamine, 10 mM glucose) for 30-60 min at 37°C in a non-CO_2_ incubator. The Glycolytic Rate Assay was performed on an Agilent Seahorse XFe96 analyzer with an acute insulin injection (100 nM final concentration). Manufacturer-recommended injection volumes were used for each assay. Final well concentrations for assay compounds are as follows: 0.5 µM rotenone/antimycin A (Rot/Ant A) and 50 mM 2DG. Data were normalized to total protein (µg) and as indicated in the figure legend. Data were analyzed using Agilent Wave software.

### Assessment of Lipid Synthesis by Thin Layer Chromatography

Adipocytes were serum-starved for 3-4 h in DMEM/0.5% BSA followed by an additional 4 h incubation with 5 µCi/well [1,2-^14^C]-acetic acid, with or without 100 nM insulin, in DMEM/0.5% BSA supplemented with 50 µM sodium acetate. Following incubation, the medium was aspirated and lipids were extracted by isopropanol for at least 24 h. Proteins from each sample were harvested using 300 µL 0.1 NaOH on a plate rocker for 30 min at RT and protein concentration was determined by the Lowry method^54^. Extracted lipids were dried under N_2_ at 50°C and resuspended in 1:1 MeOH:CHCl_3_ containing known lipid standards (cholesteryl ester [50 µg], triglyceride mixture [150 µg], oleic acid [25 µg], and cholesterol [100 µg]). Resuspended lipids from each sample were spotted on high-performance thin-layer chromatography silica plates (Millipore Sigma, Burlington, MA, USA) and separated using a solvent system containing hexane:isopropyl ether:acetic acid (65:35:2 [vol/vol/vol]). Lipid species were visualized by exposure to iodine, lipid-containing bands were scraped, and the incorporation of ^14^C radiolabel in each lipid species was measured in a liquid scintillation counter (Beckman Coulter). Data were normalized to total protein and as indicated in the figure legend.

### Insulin-Stimulated Expression of Lipid Metabolism Genes

Adipocytes were incubated in complete medium (DMEM/F12, 5% FBS, 1% P/S) in the absence of insulin for 3-4 h followed by incubation with or without insulin (100 nM) for 2 h. RNA isolation, reverse transcriptase PCR, and quantitative real-time PCR were all performed as previously described above.

### Data Normalization and Statistical Analyses

Data were normalized to WT or unstimulated controls as a fold change (normalized value = 1) or percentage (normalized value = 100%). Some raw data were initially normalized to total protein as indicated in figure legends. Statistical analyses were performed using GraphPad Prism 9.5.1 (San Diego, CA, USA). Unpaired Student’s t-tests were used to analyze two independent groups. One-way ANOVA with Tukey’s post hoc tests was used to compare three or more independent groups. For experiments with two or more independent variables, two-way ANOVA with Tukey’s post hoc tests was used. Raw or normalized data was presented as mean ± SD, where *p≤0.05, **p≤0.01, ***p≤0.001, and ****p≤0.0001.

## Abbreviations

2DG: 2-deoxy-D-glucose
ATP: adenosine triphosphate
CoA: coenzyme A
DGAT: diacylglycerol O-acetyltransferase 1
DMEM: Dulbecco’s Modified Eagle Medium
ECAR: extracellular acidification
EMSC: ear mesenchymal stem cell
FASN: fatty acid synthase
GLUT4: glucose transporter 4
glycoPER: glycolytic proton efflux rate
HDL: high-density lipoprotein
HFD: high-fat diet
KRPH: Krebs-Ringer phosphate HEPES
INSR: insulin receptor
LCAD: long-chain acyl-CoA dehydrogenase
ORO: Oil Red O
OXPHOS: oxidative phosphorylation
PCPE2: procollagen C-endopeptidase enhancer 2
PPARγ: peroxisome proliferator-activator receptor γ
Rot/Ant A: rotenone/antimycin A
SR-BI: scavenger receptor class B type I

## Author Contributions

D.A.K., Y.C., D.S., M.S.T., and M.T. conceived and designed the research. D.A.K., J.C., and Y.C. performed the research and acquired the data. D.A.K., Y.C., and D.S. analyzed and interpreted the data. All authors were involved in drafting and revising the manuscript.

## Acknowledgments

The authors would like to thank members of the Sahoo Laboratory, Hayley Powers, Gage Stuttgen, Jordan Bobek, Emma Tillison, and Renee Penoske for their critical review of this article. Further thanks are given to Kay Nicholson for technical assistance. This work was funded by the National Institutes of Health (NIH) grants F31HL149161 (D.A.K), R01HL138907 (D.S. & M.S.T.), R01HL58012 (D.S.), MCW Startup Funding (Y.C.), and was supported in part by the Redox and Bioenergetics Shared Resource (RBSR) at the Medical College of Wisconsin Cancer Center.

## Data Availability Statement

The data that support the findings of this study can be obtained from the corresponding author upon reasonable request.

## Figure Legends

**Supplemental Figure 1.**
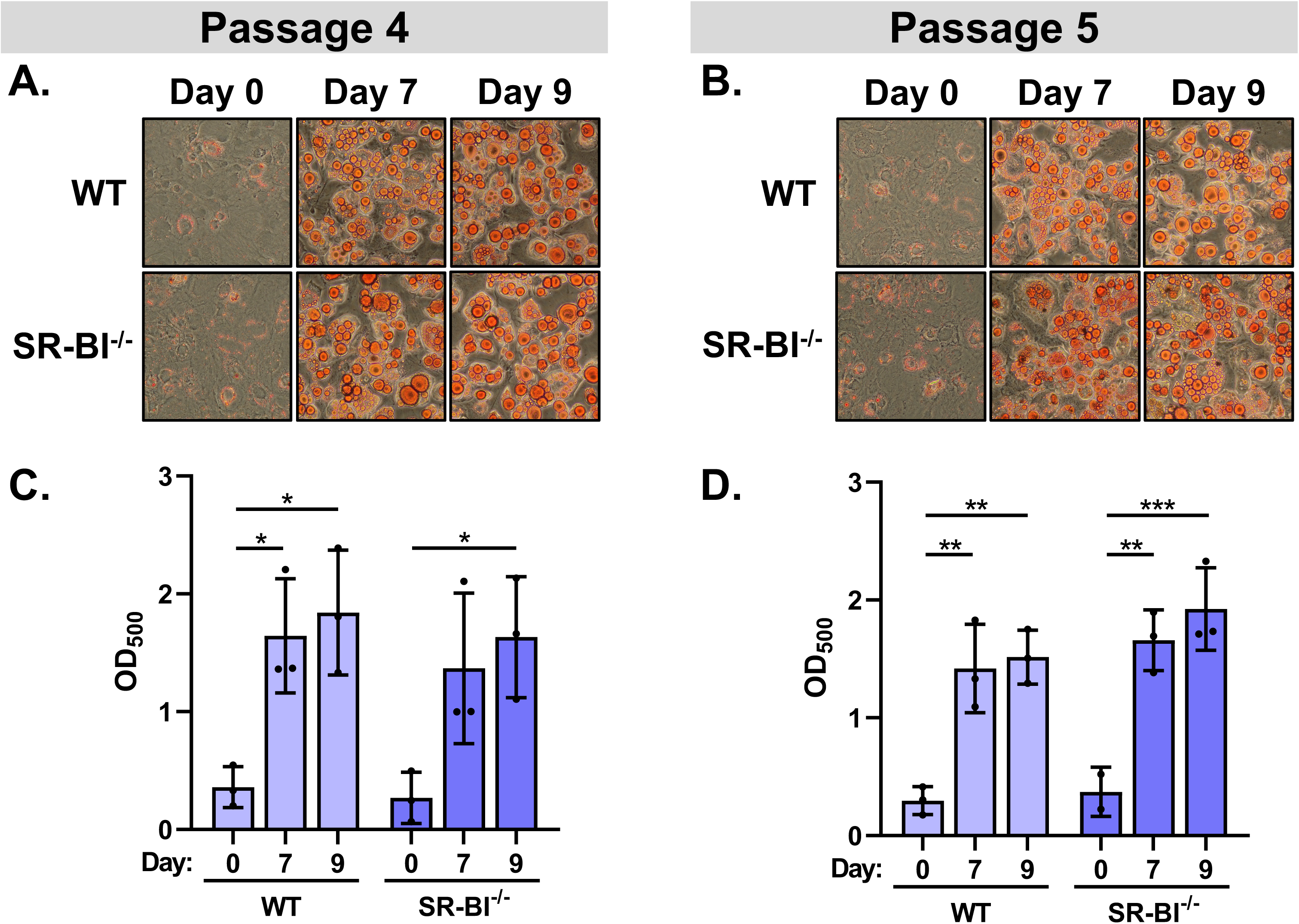
EMSCs at passages 4 and 5 can differentiate into adipocyte-like cells. EMSCs from WT and SR-BI^-/-^ mice at **(A)** passage 4 and **(B)** passage 5 were stained with ORO at days 0, 7, and 9 post-differentiation and imaged at 20X magnification. Representative images were selected (n=2-3/day/genotype). **(C)** ORO staining in passage 4 and **(D)** passage 5 was quantified by extracting the dye with isopropanol and measuring the absorbance at 500 nm. Data presented as mean ± SD (n=2-3, two-way ANOVA, Tukey’s post hoc [*p≤0.05, **p≤0.01, ***p≤0.001]).

**Supplemental Figure 2.**
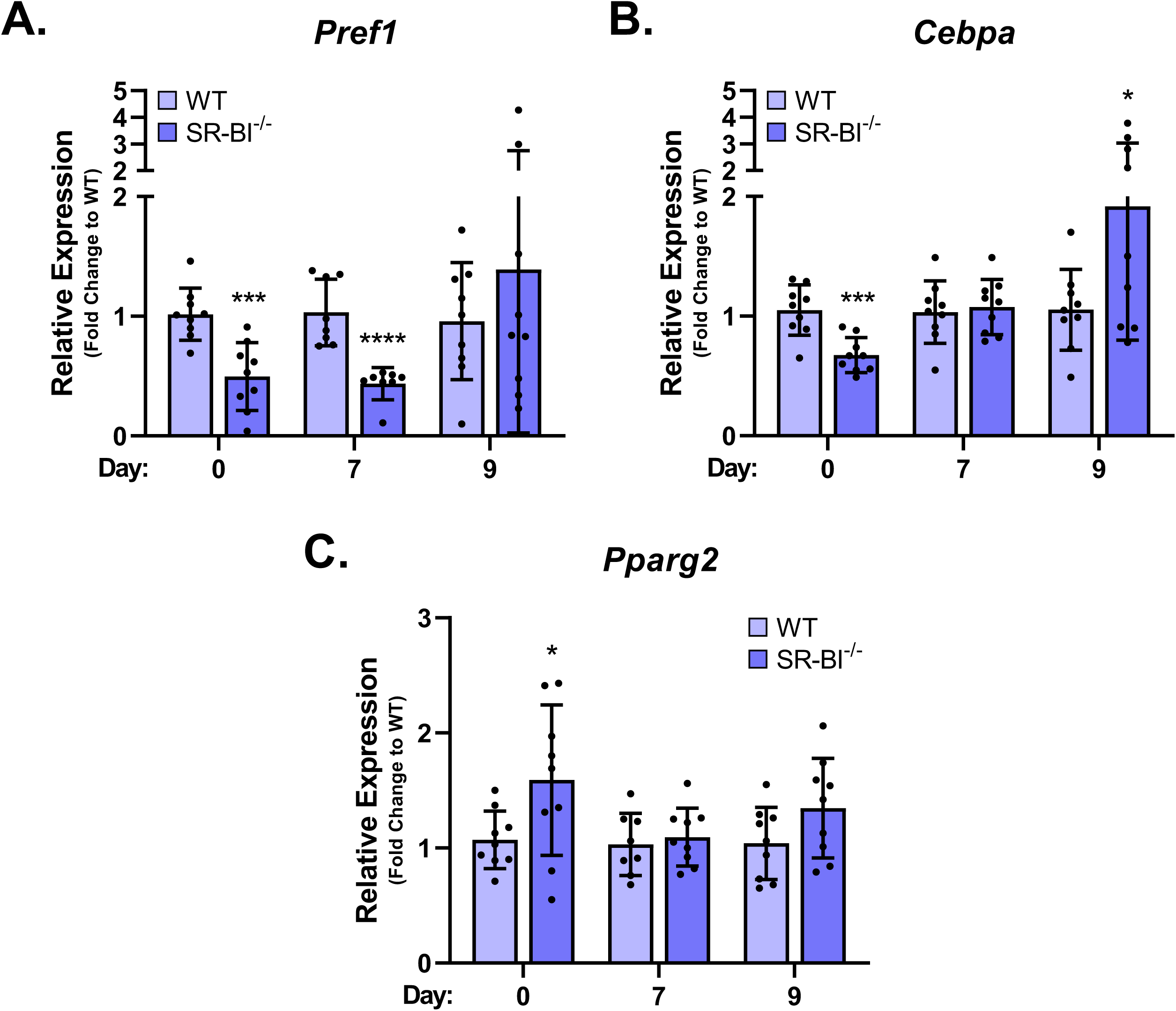
SR-BI may be important for the early stages of adipocyte differentiation. Data from Figure 2A-B were reanalyzed to measure gene expression of preadipocyte marker **(A)** *Pref1* (Pref-1) and adipocyte markers **(B)** *Cebpa* (C/EBPα) and **(C)** *Pparg2* (PPARγ2) relative to WT at days 0, 7, and 9 post-differentiation by the 2^-ΔΔCt^ method using *Rplp0* (60S ribosomal subunit) as a housekeeping gene. Data were then subsequently normalized as a fold change to WT (WT = 1) and presented as mean ± SD (n=3 independent experiments performed in duplicate or triplicate, unpaired Student’s t-test [*p≤0.05, ***p≤0.001, ****p≤0.0001]).

